# SARS-CoV-2 B.1.1.7 and B.1.351 Spike variants bind human ACE2 with increased affinity

**DOI:** 10.1101/2021.02.22.432359

**Authors:** Muthukumar Ramanathan, Ian D. Ferguson, Weili Miao, Paul A. Khavari

## Abstract

SARS-CoV2 being highly infectious has been particularly effective in causing widespread infection globally and more variants of SARS-CoV2 are constantly being reported with increased genomic surveillance. In particular, the focus is on mutations of Spike protein, which binds human ACE2 protein enabling SARS-CoV2 entry and infection. Here we present a rapid experimental method leveraging the speed and flexibility of Mircoscale Thermopheresis (MST) to characterize the interaction between Spike Receptor Binding Domain (RBD) and human ACE2 protein. The B.1.351 variant harboring three mutations, (E484K, N501Y, and K417N) binds the ACE2 at nearly five-fold greater affinity than the original SARS-COV-2 RBD. We also find that the B.1.1.7 variant, binds two-fold more tightly to ACE2 than the SARS-COV-2 RBD.

---

Genomic surveillance efforts have uncovered SARS-CoV-2 variants with mutations in the viral Spike glycoprotein, which binds the human ACE2 receptor to facilitate viral entry^**1**^. Such variants represent a public health challenge during the COVID-19 pandemic because they increase viral transmission and disease severity.^2^ The B.1.351 variant first identified in South Africa has 3 notable mutations in the Spike Receptor-Binding Domain (**RBD**), namely K417N, E484K and N501Y^**3**^ while the B.1.1.7 variant first identified in the UK carries the N501Y mutation (**Fig. S1–S3**). B.1.351 is of particular concern for its potential resistance to antibodies elicited by prior SARS-CoV-2 infection and vaccination ^**4**^.

Several mechanisms may account for increased variant transmissibility, such as increased Spike protein density, greater furin cleavage accessibility, and enhanced Spike protein binding affinity for the ACE2 receptor^5^. To test the latter, binding of purified recombinant B.1.351 and B.1.1.7 RBD was compared to the Hu-1 RBD originally identified in Wuhan (SCoV2) using microscale thermophoresis (**MST**). The B.1.1.7 RBD bound ACE2 with 1.98-fold greater affinity than the SCoV2 RBD (*K_d_* 203.7±57.1 nM, vs 402.5±112.1 nM) (**Fig 1a**). The B.1.351 RBD bound ACE2 at 4.62-fold greater affinity the SCoV2 RBD (*K_d_* 87.6±25.5 nM, vs 402.5±112.1 nM) (**Fig. 1b**). These data are consistent with a model in which variant Spike proteins mediate increased transmissibility of the B.1.1.7 and B.1.351 variants, at least in part, by enhancing ACE2 binding affinity.

**Fig. 1.**
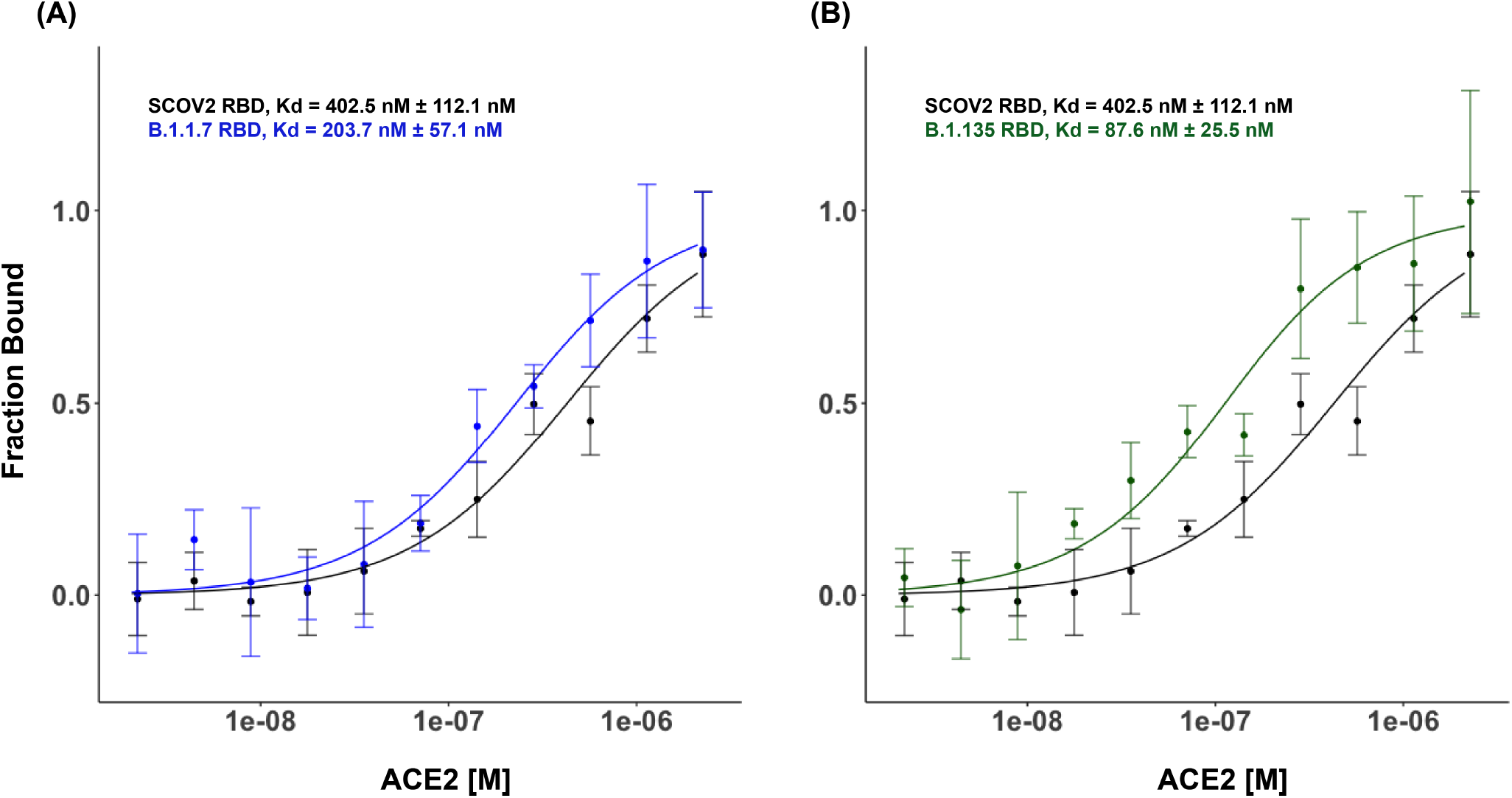
Binding of SARS-CoV-2 Spike RBD with Human ACE2. Microscale thermophoresis with 100nM labeled Spike RBD (**A**) Binding curve of SCoV2 and B.1.1.7 RBD with a titration series of human ACE2 (n=3 series of independent replicates) (**B**) Binding curve of SCoV2 and B.1.351 RBD with a titration series of human ACE2 (n=3 series of independent replicates).

These findings suggest that acquisition of enhanced affinity of Spike proteins for the human ACE2 receptor may be a convergent feature of more transmissible SARS-CoV-2 variants arising in multiple geographic regions and indicate that MST provides a rapid way to biochemically assess such changes.

## Materials and Methods

### Recombinant Proteins

Human ACE2 protein was purchased from Sino Biological (SinoBiological, catalog#10108-H05H) and re-suspended in PBS-T to obtain 4.54uM stock concentration. Recombinant Spike Receptor Binding Domains were ordered from BEI resources. Protein quality was determined using a Tycho NT.6 (**Figures S4 & S5**) (NanoTemper Technologies). Human hnRNPC was purchased from Abnova (catalog# H00003183P01S.)

### Microscale Thermophoresis

Spike RBD proteins were labeled using Monolith His-Tag Labeling Kit RED-tris-NTA (NanoTemper Technologies) following manufacturer protocol at 2:1 protein to dye ratio. Briefly, 100 uL of 200nM Spike RBD (Hu-1 or B.1.1.7 or B. 1.351 RBD) was mixed with 100 uL of 100nM dye for 30 minutes at room temperature, followed by centrifugation for 10 minutes at 15000g at 4C. A series of sixteen 1: 1 dilutions of hACE2 were prepared, with 2.27uM the highest concentration. Each dilution of hACE2 was mixed with one volume of Spike labeling reaction mix prior to incubation for 5 minutes at room temperature. Mixed samples were loaded into Monolith NT.115 Capillaries (NanoTemper Technologies). As a specificity control, hnRNPC was utilized instead of hACE2; MST showed no binding (**Figure S6**). MST was performed using a Monolith NT.115 instrument (NanoTemper Technologies) at room temperature, with 60-80% excitation power and Medium MST power.

## Acknowledgements

The following reagent was produced under HHSN272201400008C and obtained through BEI Resources, NIAID, NIH: Spike Glycoprotein Receptor Binding Domain (RBD) from SARS-Related Coronavirus 2, Hu-1 with C-Terminal Histidine Tag, Recombinant from HEK293F Cells, NR-52366. The following reagents were obtained through BEI Resources, NIAID, NIH: Spike Glycoprotein Receptor Binding Domain (RBD) from SARS-Related Coronavirus 2, South Africa Variant (NR-54005) and United Kingdom Variant (NR-54004) with C-Terminal Histidine Tag, Recombinant from HEK293 Cells. This work was supported by the USVA Office of Research and Development I01BX00140908 (P.A.K.), NIH CA142635, AR45192, AR076965 and HG007919 (P.A.K).

---

**Figure S1:**
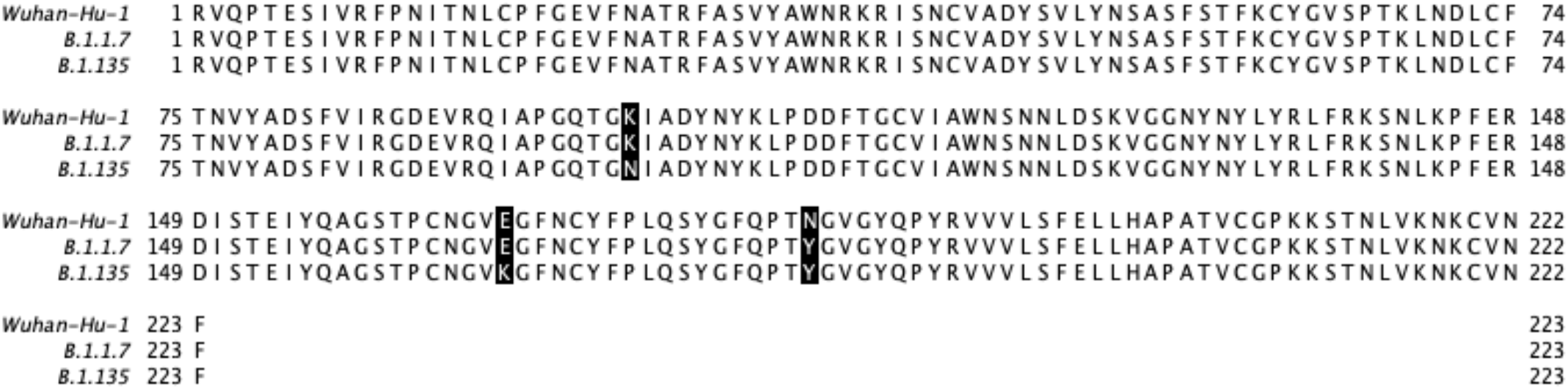
Sequence and alignment of RBD proteins assayed. The sequence of the parental Hu-1 strain is used as SCoV2 RBD sequence. Mutations seen in B.1.1.7 and B.1.351 are highlighted.

**Figure S2:**
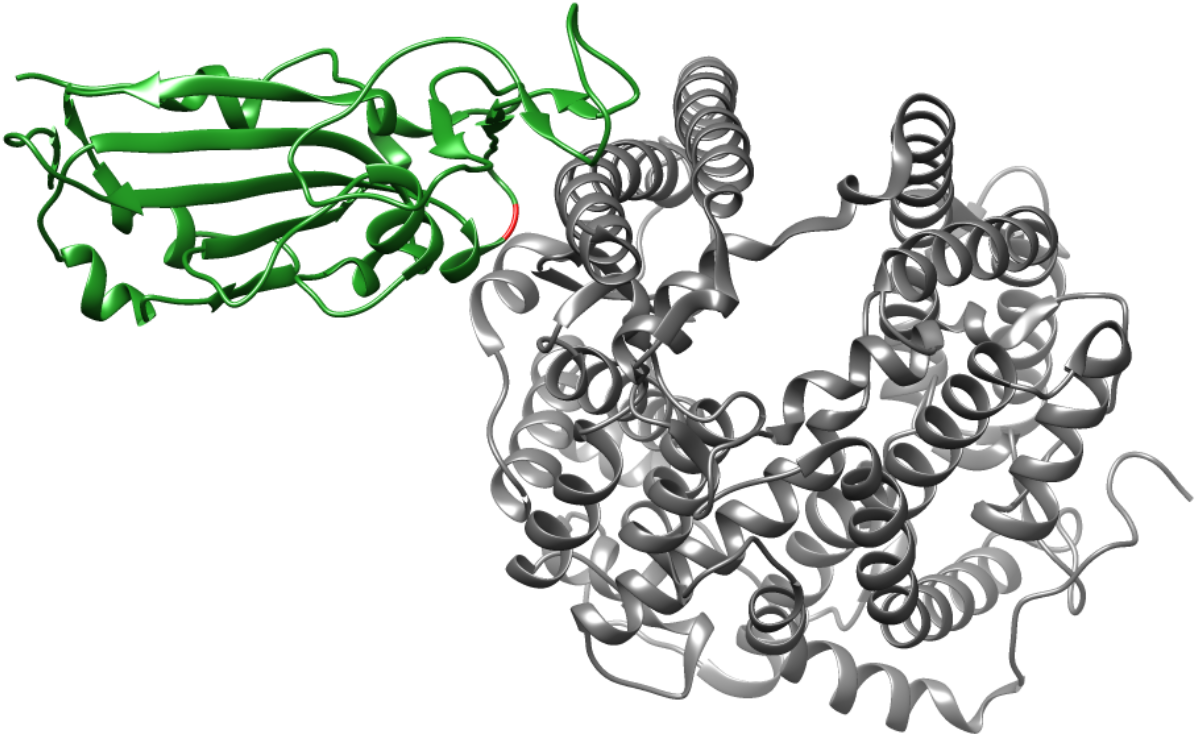
Structure of the SARS-CoV-2 RBD (PDB: 6M0J) with B.1.1.7 mutation (N501Y) on RBD colored red. Coloring schema: ACE2 (grey), receptor-binding domain (green)

**Figure S3:**
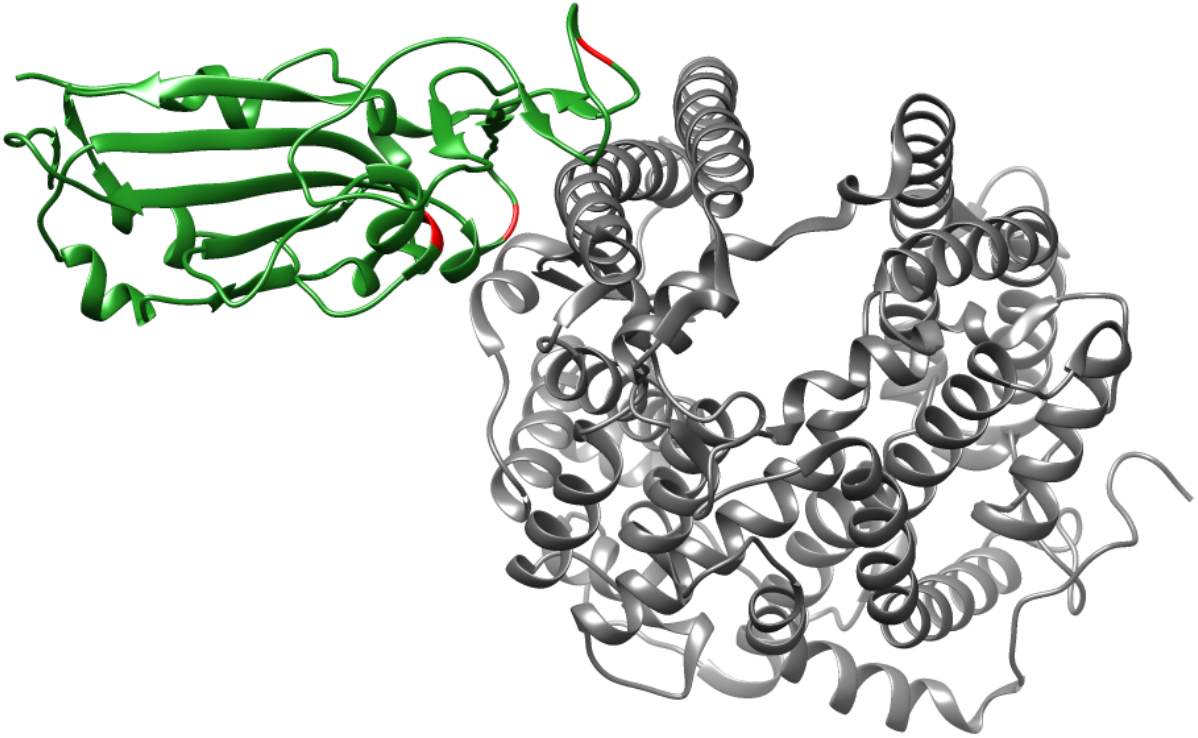
Structure of the SARS-CoV-2 RBD (PDB: 6M0J) with B.1.351 mutation (K417N, E484K, N501Y) on RBD colored red. Coloring schema: ACE2 (grey), receptor-binding domain (green)

**Figure S4:**
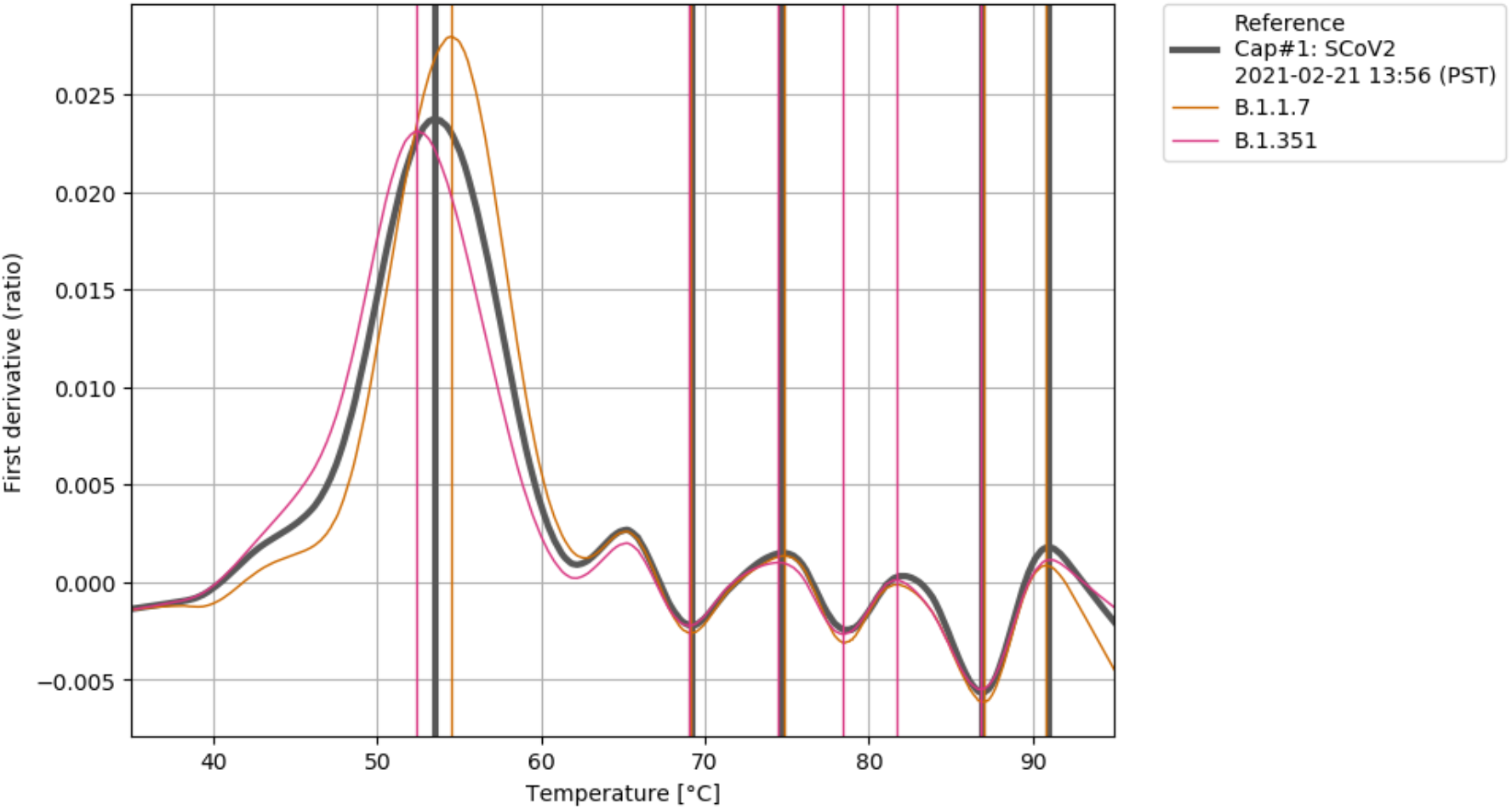
Protein quality of RBD proteins assayed. Tycho NT. 6 (Nanotemper) was used to measure changes in intrinsic fluorescence at 330 nm and 350 nm with increasing temperature. The fluorescence signals were plotted as a first derivative of the 350 nm/330 nm ratio.

**Figure S5:**
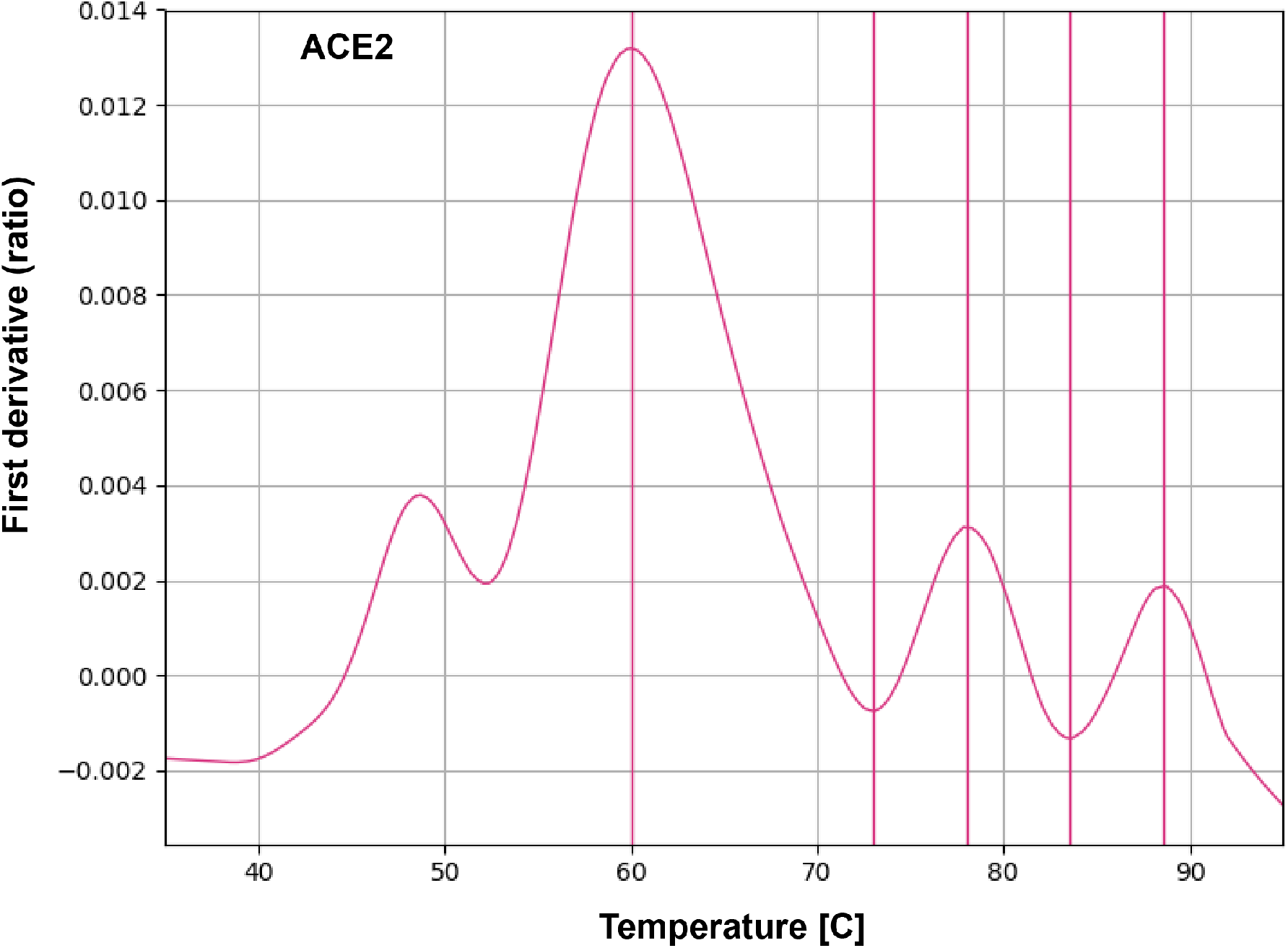
Protein quality of ACE2. Tycho NT. 6 (Nanotemper) was used to measure changes in intrinsic fluorescence at 330 nm and 350 nm with increasing temperature. The fluorescence signals were plotted as a first derivative of the 350 nm/330 nm ratio.

**Figure S6:**
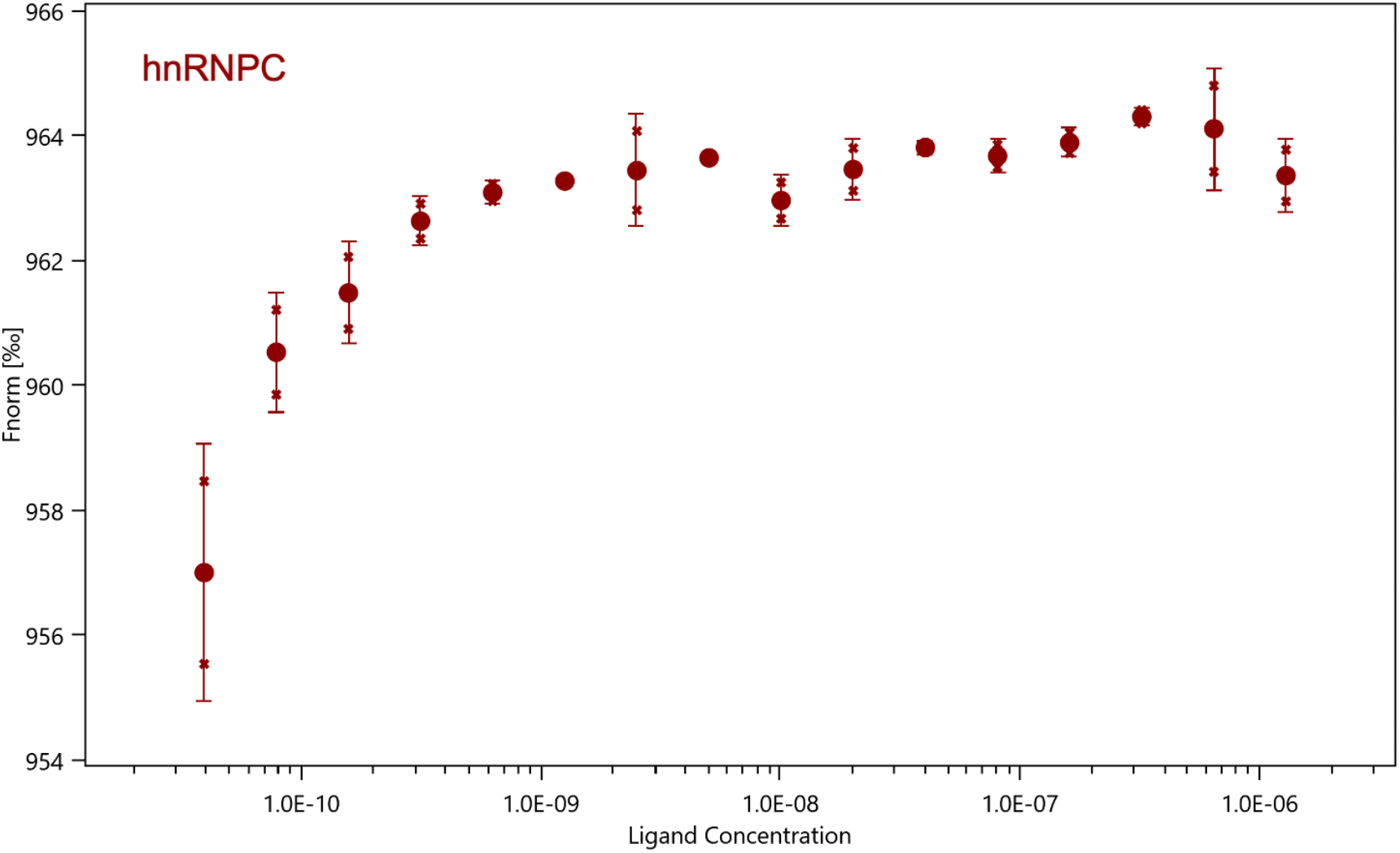
Microscale thermophoresis with 100nM labeled Spike RBD. SCoV2 and hnRNPC with a titration series of human hnRNPC protein shows no binding, confirming SCoV2 does not interact non-specifically (n=2 series of independent replicates).

